# Skin Hydration By Natural Moisturizing Factors, A Story Of H-Bond Networking

**DOI:** 10.1101/2024.10.16.618184

**Authors:** Marving Martin, Benjamin Chantemargue, Patrick Trouillas

## Abstract

Dry skin is a common condition experienced by many. Besides being particularly present during the cold season, various diseases exist all year round, leading to localized xerosis. To prevent it, the skin is provided with natural moisturizing factors (NMFs). They are small amino acids or derivatives found in the outermost layer of the skin, the *stratum corneum* (SC). They are often claimed to be highly efficient humectants, increasing the water content to maintain the fluidity of the skin. However, alternative mechanisms have been proposed, suggesting that NMFs themselves may act as lipid mobility amplifiers. This work aims at investigating the role of three NMFs, namely urea (URE), glycerol (GLY) and urocanic acid / urocanate (UCA/UCO) in SC *in silico* models, considering two different levels of humidity. Molecular dynamic simulations showed an increase in the diffusion of different lipid components, mainly free fatty acids (FFAs) and ceramides acyl chain moieties, in the presence of either high water content or NMFs. The membrane properties were modified, as seen by an increased thickness and a greater lateral stiffness. All NMFs exhibited a similar impact, whereas UCA revealed slight differences according to its charged state. By studying NMF-water intermolecular interactions, we highlighted the role of NMF as a regulator of membrane perturbations, while insuring membrane fluidity. This role allows NMFs to prevent destabilization of the skin membrane in the presence of high-water content. This study, performed at an atomistic resolution, highlighted a strong H-bond network between lipids, involving mainly ceramides but also all other components. This network can be modified in the presence of high-water concentration or NMFs, resulting in modifications of membrane properties, rationalizing hydration effects.

## Introduction

Every living organism is provided with a biological interface that maintains its integrity and protects it from the surrounding environmental aggressions. It is a key structure for the organism to evolve and adapt to the outside world threats (*e.g.,* strong climate variations, pollution, mechanic impacts or sun light).^1^ For humans, this interface is the skin, a complex multilayered and multiscale structure, composed of the epidermis, the dermis, and the hypodermis.^2^ The skin epidermis is the outer layer, in other words, the first protective barrier of the body. A major structuration of this layer is the *stratum corneum* (SC). It is a brick-and- mortar architecture, where the bricks are corneocyte cells, and the mortar is a multilamellar lipid stack. The SC is soft, relatively pliable, and capable of tolerating physical constraints or varying osmotic conditions, as long as it stays hydrated and moisturized. The corneocytes are flattened and rigid dead keratinocyte cells, containing small hygroscopic molecules named natural moisturizing factors (NMFs).^3^ NMFs bind and retain water molecules from the ambient atmosphere into the corneocytes’ lumen, in turn maintaining adequate skin moisturization, even at low humidity level. This allows enzymes to work under correct physiological conditions for biological processes such as desquamation. The mortar is an intercellular lipid matrix, surrounding and maintaining the corneocytes, which are stacked like in a brick wall. This lipid lattice is made of different ceramides (CER), free fatty acids (FFA), cholesterol (CHL), and water. All these ingredients are organized in a lamellar phase providing a tight, fluid, and effective barrier against transepidermal water loss (TEWL). To support the function of the SC, multiple topical creams have been developed using NMF, water, and skin lipids as the main ingredients. These formulations can be used in cosmetics but also to treat various skin pathologies including xerosis or atopic dermatitis.^4,5^ However, it is still quite unclear how NMF molecules and skin lipids interact at the molecular level when these topical formulations are applied to the skin. A few articles have shown the stabilizing effect of small polar compounds, such as NMF, on the intercellular lipid fluidity under reduced hydration conditions.^6–8^ Glycerol (GLY), urea (URE) and urocanic acid (UCA) demonstrated a water-like behavior, by increasing molecular mobility.^9^ These findings pave the way toward a better understanding of the role of NMFs and their skin lipid interplay to maintain the SC fluidity. In this publication, we aim at joining the efforts of molecular modeling and the literature knowledge to rationalize the underlying molecular events of NMF and/or water within the SC lipid matrix. To this aim, we used molecular dynamics (MD) as a powerful tool to sample all molecular interactions and motions of a series of three NMFs embedded into an *in silico* model of the SC intercellular lipid matrix. The three NMFs studied here are URE, GLY and UCA. They were inserted in a SC model of reference and at two different hydration levels. Both UCA and its deprotonated urocanate (UCO) forms were considered, as the pKa of this acid-base couple is about 3. Although the pKa value can be modulated at the membrane interface and inside the membrane, at skin pH, the negatively charged UCO form is likely to be present, possibly predominant, in the hydrated zones of the skin.

## Computational details

The all-atom skin model of reference (REF) was inspired by an early published article.^10^ The REF model is designed to reproduce cryo-EM (electron microscopy) structures^11^ of the intercellular lipid matrix obtained from near-native human skin. It is composed of fully extended (splayed) CER, FFA, CHL, and water molecules with the following distribution 90:88:88:90, respectively. Three types of ceramides sphingoid bases were used in this system, with a physiological acyl chain length distribution,^12^ namely non-hydroxylated sphingosine (NS), non-hydroxylated phytosphingosine (NP), and esterified ω-hydroxylated sphingosine (EOS). The complete composition of the REF system is given in Table 1. According to the original publication, a specific 3D spatial arrangement of the lipids is required to accurately mimic the EM structures, the water permeability and the thermotropic behavior. Hence, 75% of CHL is located close to the CER sphingoid chains, and the remaining 25% CHL located in close contact with the CER acyl chains. FFAs are only positioned close to the acyl chain moieties of CER whereas water is inserted close to the lipid polar headgroups. To construct the REF model, each molecule topology was built from scratch, using the MarvinSketch software.^13^ The REF system, containing 90 water molecules, was manually assembled in the VMD software^14^ to form the initial structure (Figure 1**.a-b**). The lipids and water were described by the CHARMM36 forcefield^15^ and the TIP3P model, respectively. The adjusted polar head dihedral parameters were collected from the publication’s supplementary data^10^. Energy minimization was performed using GROMACS 2021.^16^ The minimized structure was then heated to 305K (in the NVT ensemble) for 1ns, followed by a semi-isotropic pressurization maintained at 1 bar for 1ns (in the NPT ensemble). The output structure displayed four different layers namely: short-chain, lipid headgroup, long-chain and liquid-like layers (Figure 1**.c**). The short-chain layer is made of 75% CHL and all CER sphingoid chains. The long-chain layer contains ceramide acyl chains, FFA and 25% of CHL. The liquid-like layer is a highly disordered region containing CER EOS ester moieties. The different physical-chemical parameters of the membranes confirmed that the REF system behaves as expected. For example, the calculated overall thickness was found very similar to the value found in the original publication, at around 10 nm (see Figure 1**.c**).^10,11^ The density along the z-axis confirmed the positioning of the different lipids and the presence of water molecules, mainly in the lipid headgroup regions and a smaller amount in the liquid-like layer (see Figure 2**.a**).

**Figure 1.**
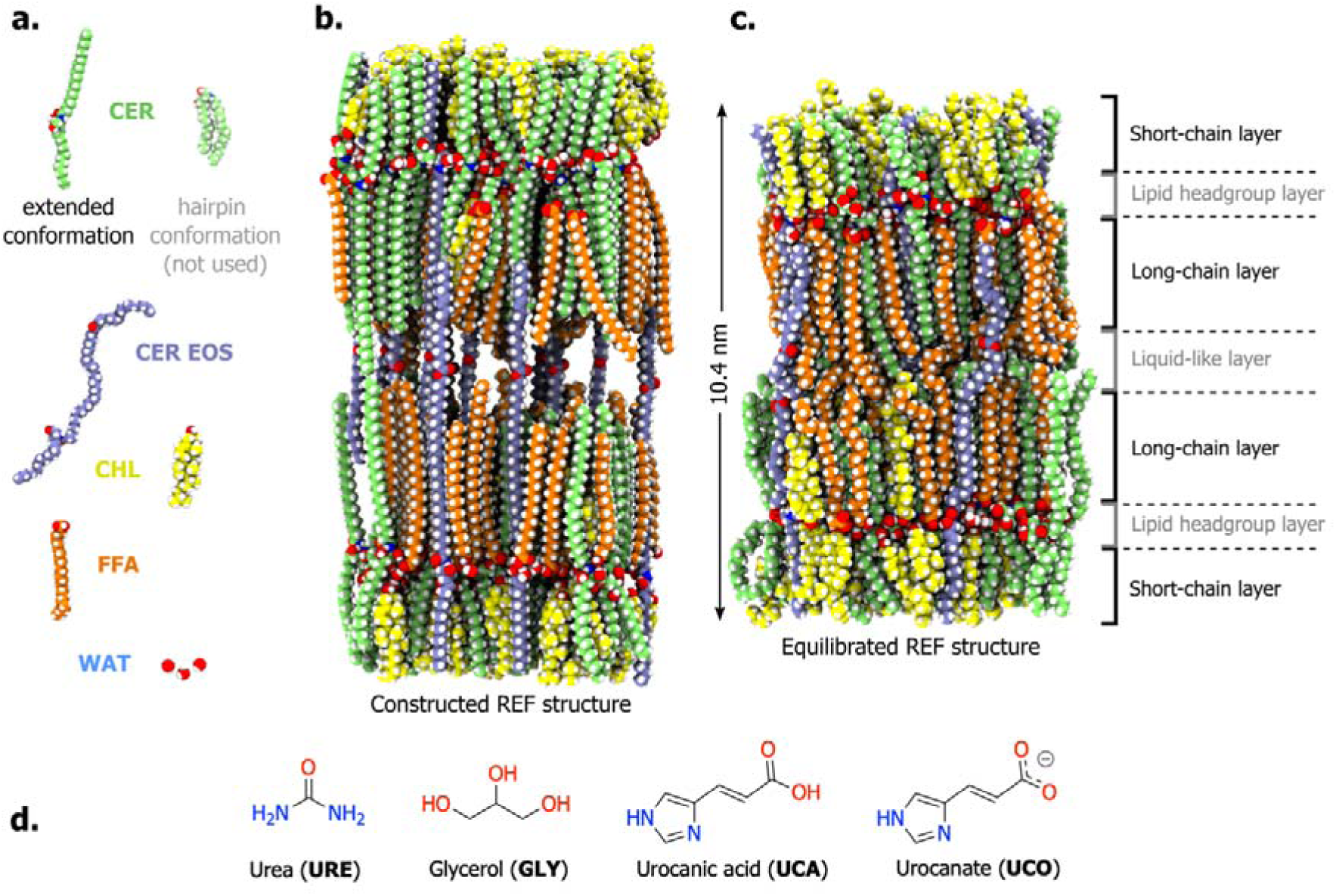
(a.) REF system composition: CER NS and NP in green, CER EOS in purple, CHL in yellow, FFA in orange and water. (b.) Initially constructed REF structures using extended CER conformation. (c.) Equilibrated REF structure revealing the organization of the different layers along the z-axis. (d) NMF chemical structures.

**Figure 2.**
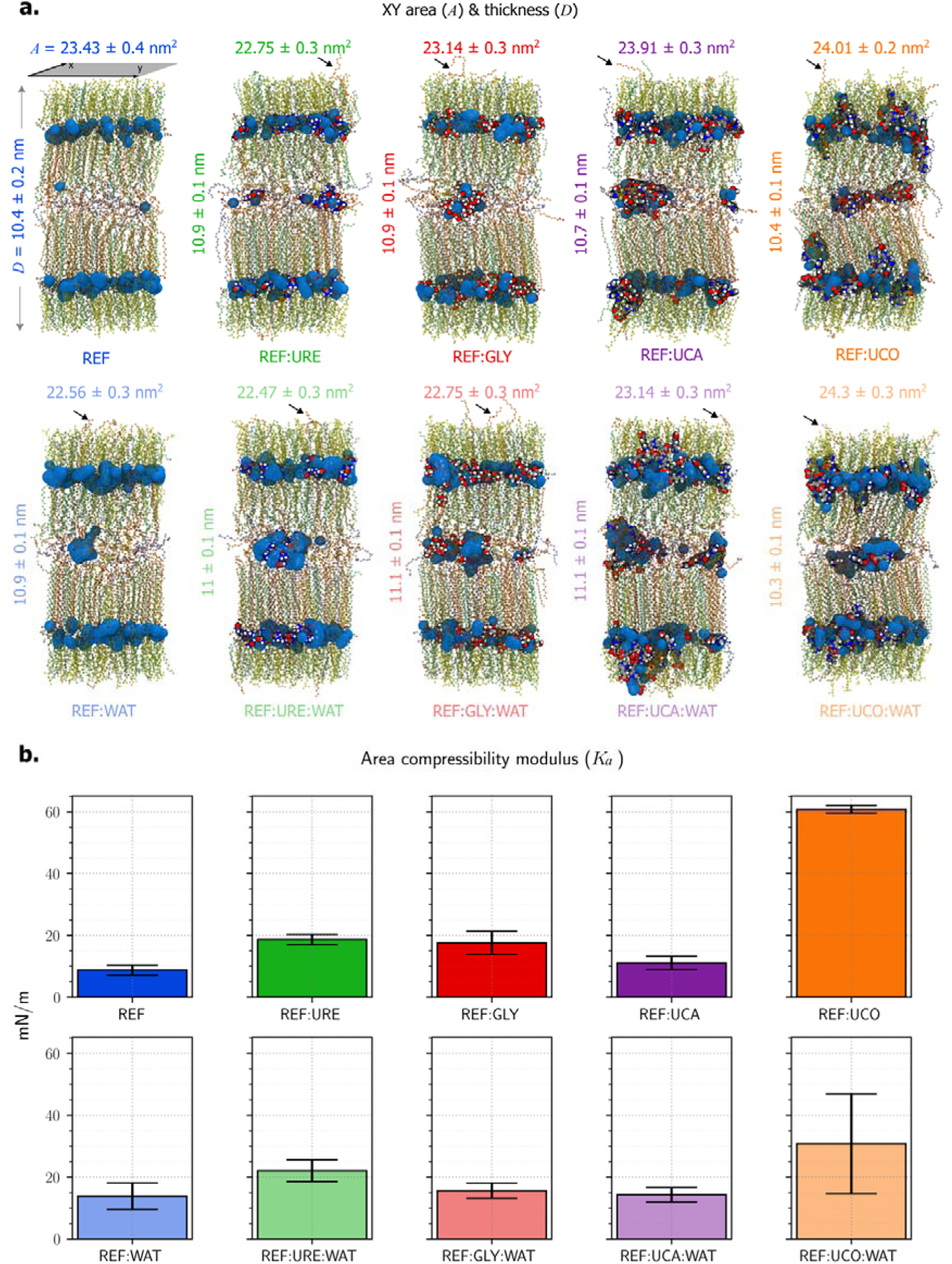
(a.) Trajectory snapshots of the different systems highlighting the position of water (blue surface) and NMF molecules in van der Waals spheres within the SC lipids (CERs in green, CER EOS in purple, FFA in orange and CHL in yellow represented as balls-and-sticks). The z-axis thickness and XY-area value of each simulation box are averaged from trajectory and replica. The arrows indicate the FFA located in the short chain layer. (b.) Trajectory average of xy-plane area compressibility modulus (*K_a_*) of each system, describing their stiffness.

**Table 1.**
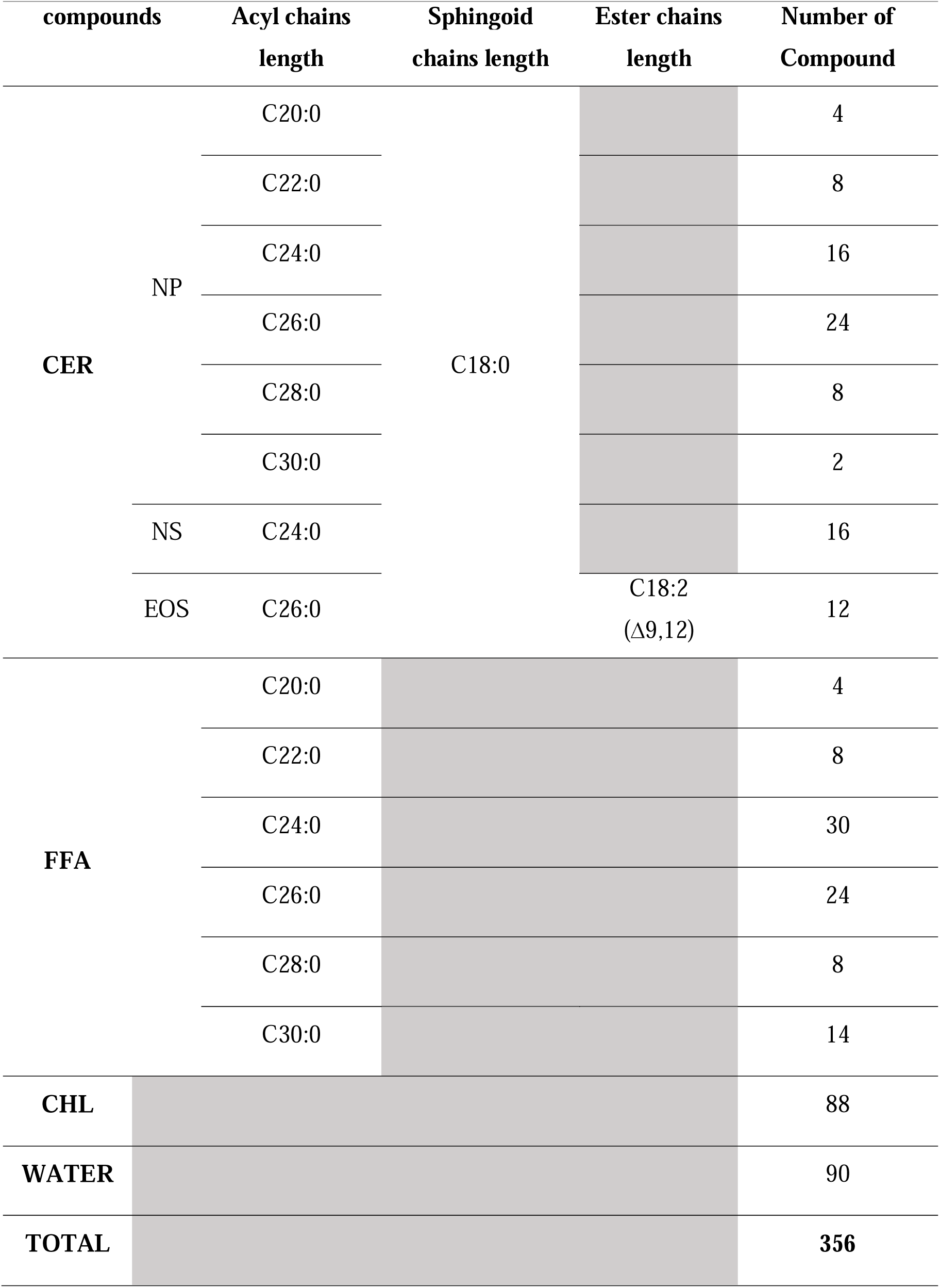
Detailed composition of the lipid matrix model of reference *i.e.*, REF system.

The production was run for 500 ns, with both electrostatic and Lennard-Jones cutoffs set at 12 Å, and the longer-range interactions were performed using the Particle-Mesh Ewald procedure. The timestep integration was set to 2 fs and the LINCS algorithm handled hydrogen bond restrain. The REF:WAT system, containing a doubled amount of water molecules, *i.e.,* 180, was created from a fully-relaxed REF model by increasing the water content close to the lipid headgroups. Both REF and REF:WAT systems were used to create eight other systems containing, each, 72 NMF (URE, GLY, UCA or UCO) molecules equally partitioned to each lipid headgroup layer and the liquid-like layer (see Figure 1**.d**). The concentration of NMF was calculated to be 20% of the number of lipids to be close to the experimental condition, namely 72 molecules.^9^ Therefore, the following eight systems were studied: four REF:NMF systems (REF:GLY, REF:URE, REF:UCA and REF:UCO) and four doubled-hydrated systems, namely REF:GLY:WAT, REF:URE:WAT, REF:UCA:WAT and REF:UCO:WAT. All NMF geometries were optimized using the Gaussian09^17^ program at the MP2/6-31G* level of theory, while the bonded parameters and charges were collected from the CGenFF webserver.^18^ UCA and UCO molecules were obtained in *trans*.^19^ The bonded parameters were automatically assigned by atom type similarity, while the charges were optimized through an extended charge increment scheme, developed by the forcefield authors.^18^ This charge-optimization method adjust partial atomic charges, by evaluating the local environment of atoms so as to capture inductive and mesomeric effects across the molecules for up to three atoms. In the end, the algorithm evaluated the charge assignment with a low penalty score for all NMF molecules, confirming the robustness of the parameters with no further optimization process.^18^ MD productions were run in triplicate, for each of the eight systems. The convergence of the simulations was monitored using area per lipids (Figure S1) and the position of the molecule along the z-axis (Figure S1, S2, S3) during the simulation time. Most of the analyses were carried out using AmberTools22 packages.^20^ The electrostatic potential was assessed with the GROMACS analysis tools, and the percentage of CER hairpin conformations (see Figure 1**.a**) was calculated with an in-house Python script. The xy-plane area compressibility modulus (*Ka*)^23^ was calculated in Figure 2**.b** using the following formula:

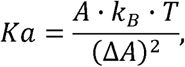

where k_B_ is Boltzmann’s constant (1.38×10^−23^□J/K), T is the temperature in K, and A is the area per lipid. Also, in order to interpret the results correctly, the impact of UCA was studied all along the membrane z-axis, whereas for UCO only the hydrated lipid headgroup layer was considered, *i.e.*, regions where this charged state is likely to exist.

## Results

### 1. Lateral stiffness and skin swelling upon hydration

When exposed to a humid environment, the SC has been described to absorb and accumulate a significant amount of water, leading to the swelling of the skin^21^. Although this swelling effect is often attributed to the swelling of corneocytes^21^, here we show that it is also partly rationalized by a behavior of the SC lipid matrix at the molecular level, by comparing the MD simulations between the REF system and the REF:WAT system. The lipid matrix thickness (*D*) increased upon water addition by 0.5 nm (see *D*_REF_ vs *D*_REF:WAT_ in Figure 2**.a**). This increase mainly comes from a swelling of the liquid-like region where both FFA and water partition in greater amounts as humidity increases (see the water blue surface in the middle of the snapshot for the REF:WAT compared to that of REF, in **Figure 2.a**). Concomitantly to the thickness increase, the xy-area (*A*) decreased by *c.a.* 0.9 nm^2^ (see *A*_REF_ *vs A*_REF:WAT_, in Figure 2**.a**), suggesting a change in the skin elasticity, as seen experimentally^22^. This can also be followed by the *Ka* (Figure 2**.b**).^23^ This parameter estimates the membrane resistance to deformation in the xy-plane. Comparison between *Ka*_REF_ and *Ka*_REF:WAT_ depicted an increased lateral stiffness of the lipid matrix model by 5 mN/m. This means that as the system swells upon hydration, the xy-plane is more resistant to compression, as reflected by the increased *Ka*. This suggests an increase of the lipid matrix lateral organization. Such an increase brings a focus on how water can reshape the physical interactions around the lipid headgroups, as detailed in the discussion section.

### 2. Location of NMFs in the SC lipid matrix and molecular impact

The three NMF molecules (URE, GLY, and UCA) are mainly located in the lipid headgroup region and fewer in the liquid-like region (**Figure 2.a** and **Figure S4** for a representative snapshot or the density profile, respectively). In these positions, they induce a similar swelling and lateral stiffness of the membrane, as that observed when the water concentration was increased (REF:WAT system). Indeed, an increase in the thickness and the *Ka* was observed (**Figure 2)**, when comparing REF to REF:URE (+0.5 nm, +10 mN/m), to REF:GLY (+0.5nm, +8 mN/m) or to REF:UCA (+0.3 nm, +2 mN/m).

The UCA molecule and its charged counterpart UCO are π-conjugated, rigid and cylinder-in-shape molecules. Their orientation with respect to the z-axis is subjected to rapid movements (**Figure S5**). This induced an increase in the xy-area (*c.a.* +0.5nm^2^) in the REF:UCA and REF:UCO systems compared to the REF system. Conversely, according to the *Ka*, the system with UCO exhibits a higher lipid matrix lateral configuration than REF as observed by a +50 mN/m increase of the lipid lateral stiffness (**Figure 2.b**). The global trend is that the NMFs behave as water does in terms of membrane properties. This is in line with a previous study showing the water-like effects of NMF on pig skin.^9,24,25^

### 3. NMFs role upon hydration

In the presence of both NMFs and water increase, the swelling and lateral stiffness effects were not additive. Namely, no further improvement in the lipid matrix lateral organization was seen, as area compressibility modulus means and standard deviation were comparable (*e.g.*, 18±2 mN/m for *Ka*_REF:URE_ vs 22±4mN/m for *Ka*_REF:URE:WAT_ in Figure 2**.b**). An increase in the swelling of the lipid matrix was observed but not equal to an addition of NMF and water effects. When comparing *D*_REF_ to *D*_REF:URE:WAT_, only a 0.6 nm thickness increase was observed (see in Figure 2**.a**), instead of an expected 1.0 nm increase in case of additive NMF and water effects (see 0.5 nm increase from *D*_REF_ to *D*_REF:URE_ and 0.5 nm increase between *D*_REF_ and *D*_REF:WAT_ in Figure 2**.a**). The same trend can be observed for the other two NMF molecules (GLY and UCA). This suggests an antagonist effect of water and NMF when they are both added to the physiological (REF) system. As water content increases, the literature has shown a high risk of skin lipid matrix disorganization.^9^ This was rationalized by the formation of water pools at the molecular level.^24,26^ The early-stage formation of such water pools can be observed here from the MD simulations, at the liquid-like layer of all systems containing an increased water content (see water aggregation in this region for REF:WAT, REF:NMF and REF:NMF:WAT in Figure 2**.a**). Conversely to all other systems, REF:UCO was the only system for which water addition in REF:UCO:WAT increased the xy-area while maintaining its thickness constant (see 24±0.2 nm^2^ and 60±2mN/m for REF:UCO vs 24.3±0.3 nm^2^ and 31±17mN/m for REF:UCO:WAT in Figure 2). This behavior is rationalized in terms of molecular interactions in the Discussion section.

### 4. Lipid mobility

As a result of a high hydration level (REF:WAT), a few FFA molecules migrate from the long-chain to the short-chain layers (see arrows in Figure 2**.a**). This migration suggested an increase in lipid mobility upon hydration. Figure 3 presents three graphics helping to monitor the lipid dynamics and interlayer crossing events. The diffusion coefficient of FFA increased (*i.e.,* by + 5.10^8^ cm^2^/s, see Figure 3**.a**). The same applies to CHL but to a lower extent than FFA (see **Figure S6**). This mobility increase can also be indirectly seen by a lower FFA order parameter (**Figure S7**). The increased diffusion rationalizes the FFA infiltration into the short-chain layers, as observed by the presence of FFA in this region in the REF:WAT system (Figure 3**.b**). The CER lipids also indirectly exhibited a greater mobility in the REF:WAT with respect to the REF system. This is partially attributed to a lower order parameter of their acyl chain moieties (**Figure S7**) but also and mainly due to changes in their conformation, namely from splayed-to-hairpin, with a hairpin conformation increase by more than 10% (Figure 3**.c**). This change is consistent with the changes in CER conformation proposed by Norlén *et. al.* during the formation of the skin barrier in the SC.^27^ In the latter publication, the authors explained that lipid dehydration induced molecular stretch-out of the CER from the deeper to the upper layer of the viable epidermis, resulting in a hairpin-to-splayed conformational change. Here, we show the reverse process splayed-to-hairpin upon hydration, highlighting how crucial is the control of skin hydration, a matter of importance that must be seen at the molecular level.

**Figure 3.**
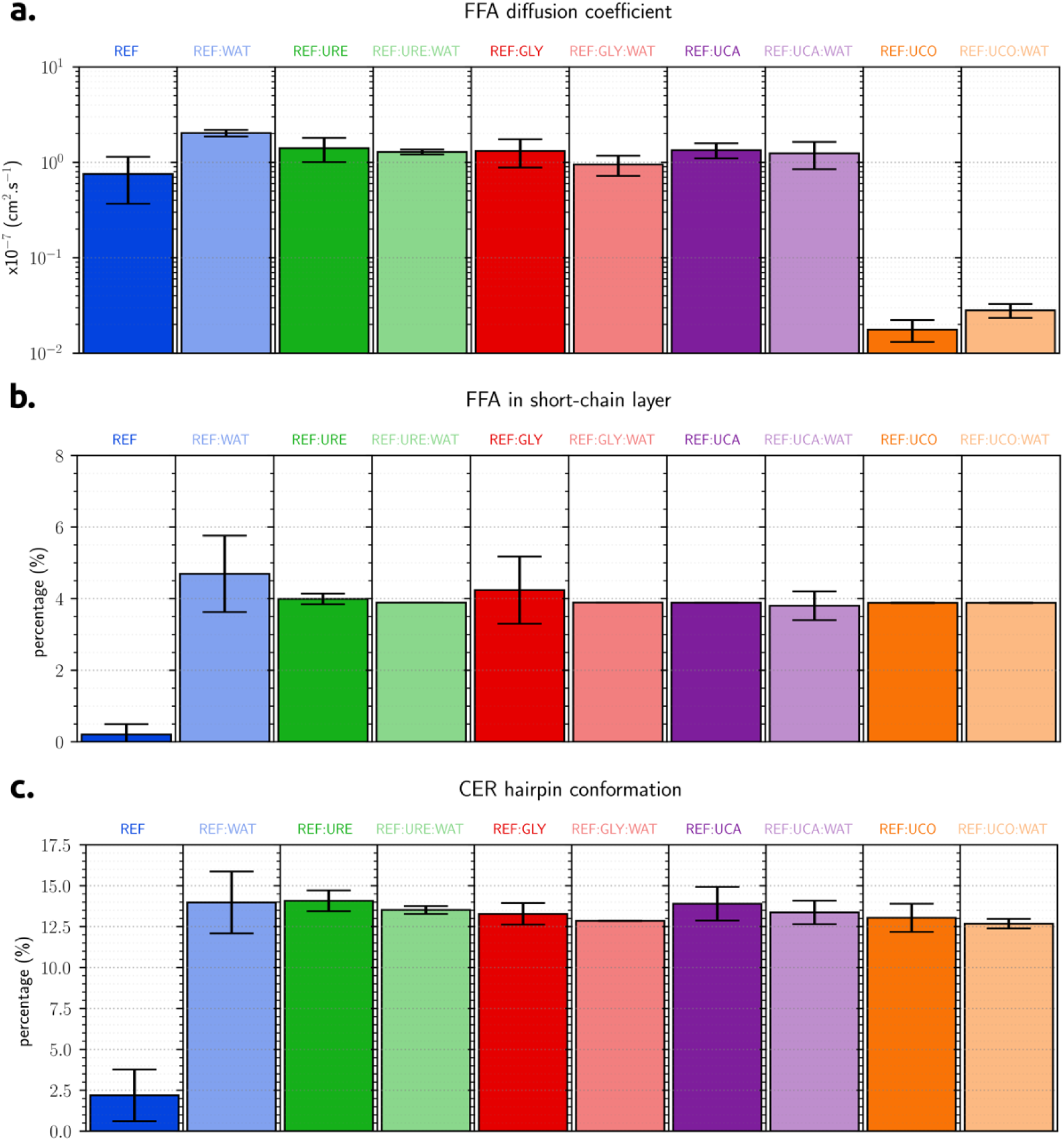
(a.) Diffusion coefficients of free fatty acid (FFA). (b.) Percentage of FFA found in the short chain layers. (c.) Percentage of ceramides (CER) hairpin conformation found among ceramides.

Interestingly, when incorporating the NMFs (URE, UCA, and GLY) into the lipid matrix, similar effects were observed, namely more mobile lipids with increased FFA diffusion coefficients (Figure 3**.a**), so short chain layer infiltration by FFA molecules (arrows in Figure 2**.a** and Figure 3**.b**), and presence of CER hairpin conformation (Figure 3**.c**). Conversely to the other three NMFs, the presence of UCO (REF:UCO system) dramatically reduced the FFA diffusion coefficients compared to the REF system.

## Discussion: rationalization of underlying interactions

Behind the structural modifications of the lipid matrix of the SC lies a complex H-bond interaction network between all molecules. Figure 4 summarizes the characterization of this network. In the REF system, CER lipids exhibit the greatest capacity for H-bond interactions (the average heteromolecular H-bonding interaction involving CER, *i.e.,* CER-X, being 106%, in Figure 4**.a**). It is worth mentioning that these lipids maintain at least one H-bond with another molecule all along the simulation. This makes a strong interaction, maintaining each component at its respective z-position, preventing lamellar disorganization. As such, CER is a keystone molecule of the lipid matrix organization, and therefore of the skin barrier function, as already suggested. ^28^ The same interaction was suggested in an earlier article using infrared spectroscopy.^29^ Later on, in MD simulations of the lipid matrix using CER in hairpin model, others highlighted H-bonds between CER and the amide and carbonyl groups as key in these non-covalent interactions^30^. CHL and FFA are less involved in the H-bond network (the averaged heteromolecular H-bond being 55% and 59%, respectively, in the REF system, the lowest seen in Figure 4**.a**). As a consequence, the diffusion coefficients of FFA and CHL are higher than those of CER (**Figure S6**).

**Figure 4.**
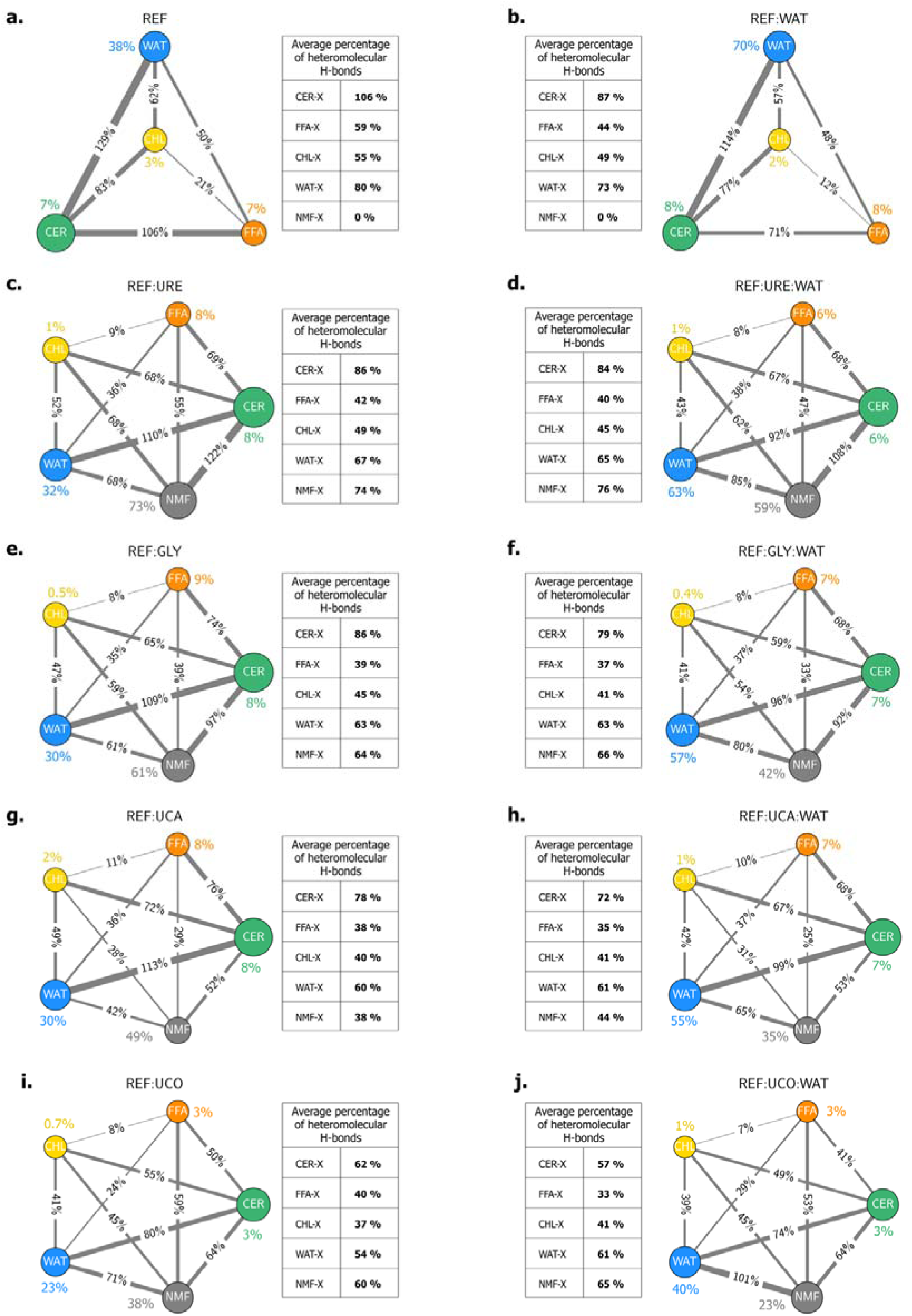
Percentage of H-bonding interactions established between molecules within each system from (a.) to (j.). An H-bond is defined as when the distance between the donor and acceptor is less than 3.5 Å and the angle is greater than 120°. The size of each circle is proportional to the amount of H-bonding interaction established by the molecule. The colored percentage value refers to self-interaction. The sum of heteromolecular H-bonds concerning each compound is calculated, then averaged, and reported in the tables. Percentages over 100% mean more than one H-bond per molecule pair are formed.

In the highly hydrated system (REF:WAT), CER lipids are still the most involved in the H-bond structuring network (Figure 4**.b**). However, the number of H-bonds decreased compared to the REF system (*c.a.* -19%, -15%, -6%, -7% for CER, FFA, CHL and WAT, respectively). This leads to a decrease in the intermolecular friction in this system, allowing higher diffusion values than in the REF system, as indicated by the disordered CER acyl chains and FFAs (**Figure S7**). Another consequence is that water molecules diffuse more (**Figure S6**) and populate more the lipid headgroup and the liquid-like layers (**Figure S4**). During these movements, water drags FFA molecules, which can also reach the liquid-like layer but also protrude to the short-chain layer. This can also be observed by a significant loss of H-bond interaction between CER and FFA (decrease from 106% to 71% in Figure 4**.a** and **b**). In the short-chain layer, the presence of FFA induces a higher order parameter of the CER sphingoid moiety (**Figure S7**). Meanwhile, in the liquid-like layer, FFA molecules are subjected to rapid motions, changing their orientation from parallel to perpendicular with respect to the z-axis. During these migration events, FFAs create voids that are quickly fulfilled by CER, where splayed-to-hairpin conformational changes can occur (see animation in **S.I**). This cascade of events modifies the lipid lamellar organization, as seen by the increased thickness and *Ka* indicating a swelling of the membrane and an increase of the lateral stiffness. Such changes are likely driven by the increase in water-water H-bonding (from 70% to 38% WAT-WAT in Figure 4**.a** and **b**).

In the presence of NMFs, namely REF:URE, REF:GLY, REF:UCA systems, the Result section reported a similar lateral stiffness, swelling, and FFA mobility effect as in the presence of high-water content (REF:WAT). This is attributed to the presence of an additional partner (the NMF) in the H-bond network, hence weakening H-bond connections between the other partners (see averaged heteromolecular H-bonds in Figure 4**.c, e, g** and **i**). As being the most bonded molecules, CER maintained a strong connection with water and NMF, while decreasing the number of connections with FFA (*e.g.*, -37% CER-FFA, -19% CER-WAT when comparing REF, in Figure 4**.a**, to REF:URE, in Figure 4**.c**). As a result, FFA molecules diffused at a higher rate in the REF:NMF systems, than in the REF system, but slightly lower than in the REF:WAT systems. This is attributed to the formation of new FFA-NMF H-bonds, as NMFs are small molecules which easily migrate in the different layers together with water, also dragging FFAs. This weakening effect of H-bonding was also observed on a similar membrane model in the presence of a series of excipients.^31^

When the hydration level increases in the presence of NMF (WAT:NMF systems), the water-NMF H-bonding connection is the major impacted observable, with +17%, +19%, +23%, from REF:URE to REF:URE:WAT (Figure 4**.c** and **d**), from REF:GLY to REF:GLY:WAT (Figure 4**.e** and **f**) and from REF:UCA to REF:UCA:WAT (Figure 4**.g** and **h**), respectively. This modification highlighted a competition between water and NMFs in the formation of H-bonds. The presence of NMFs modulates the impact of water on the lipid matrix organization and its role in the supramolecular H-bond network. FFA protrusion in the short-chain layer is then limited, preventing further swelling that could lead to lamellar disorganization. The acyl chains (*i.e.*, FFAs and acyl chain moieties of CER) appear more ordered in the REF:NMF:WAT systems than in REF:WAT.

All NMFs behaved similarly with the exception of UCO which does not induce a swelling effect despite having FFA protruding in the short-chain layer (**Figure 2.a**). This example reveals that the swelling of the lipid matrix cannot be only attributed to FFA protrusion, but it should be understood by considering all possible molecular interactions happening in the membrane. Here, the absence of swelling is attributed to the negative charge state of UCO, which shifted the electrostatic potential from positive to negative, enabling stronger interactions with its surroundings, mainly by creating salt bridges (Figure 5). These strong interactions are mainly established with water, considerably slowing down the diffusion of the two molecules. As a result, the membrane appears partially blocked in a local potential energy well.

**Figure 5.**
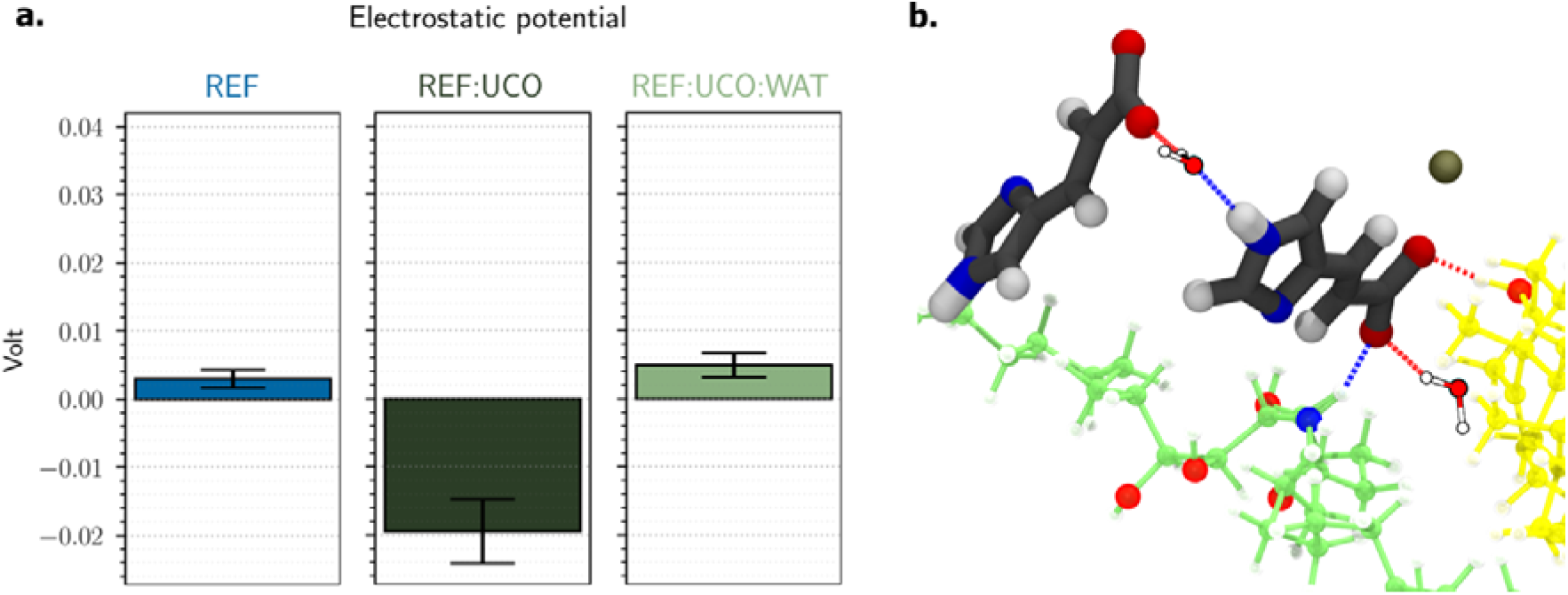
(a.) Overall electrostatic potential measured for each system. (b.) Snapshot of UCO (gray) establishing salt bridges (red) and H-bonds (blue) with CER molecule (green), CHL (yellow), sodium anion (gray ball), and water.

## Conclusion

Within a molecular level perspective, this study has confirmed the individual and mutual effects of water and NMFs when inserted in the SC intercellular lipid matrix. Under physiological conditions, CER are keystones of the supramolecular organization of the lipid matrix, by establishing a solid H-bond network with other lipids, contributing to the stability of the lipid matrix. The physical-chemical properties of this lipid matrix were monitored by following the evolutions of this H-bond network. These observations help rationalize the macroscopic impact of hydration and NMF on the skin.

When the amount of water molecules increases in the lipid matrix, they locate at the lipid polar head region. In turn, they restructure the interaction network by weakening some H-bonds, especially those involving FFAs, thereby increasing their mobility. Consequently, some FFA molecules migrate in between the layers, inducing voids and CER splayed-to-hairpin conformational switches. This increases the lateral stiffness of the lipid matrix as well as the membrane thickness or swelling. The three NMFs, URE, GLY and UCA, exhibit the same effect as that observed with high-water content, as already experimentally suggested.^9^ In presence of UCO, we did not observe a swelling of the membrane despite FFA migration in shorter chain layer. This observation showed that the swelling is not only due FFA protrusion, but to all possible molecular interactions occurring in the membrane, which can be taken in its entirety by molecular dynamic simulations. This study also highlights the beneficial interplay between water and NMFs within a large supramolecular H-bond network, as the two compounds work together through preferential interactions to enhance skin hydration and to maintain a proper lateral and lamellar organization throughout the whole membrane.

## Acknowledgments

We thank CALI (CAlcul en Limousin), Florent Di Meo and Xavier Montagutelli for computational resources as well as InSiliBio and INSERM for financial support.

