## Supporting information for "Skin Hydration By Natural Moisturizing Factors, A Story Of H-Bond Networking"

a) INSERM U1248 Pharmacology & Transplantation, Univ. Limoges, CBRS, 2 rue du prof. Descottes, F-87000 Limoges, France

b) InSiliBio, 1 avenue d’Ester, Ester Technopôle, F-87000 Limoges, France

c) RCPTM CATRIN Palacky University, Slechtitelu 27, 783 71, Olomouc, Czech Republic.

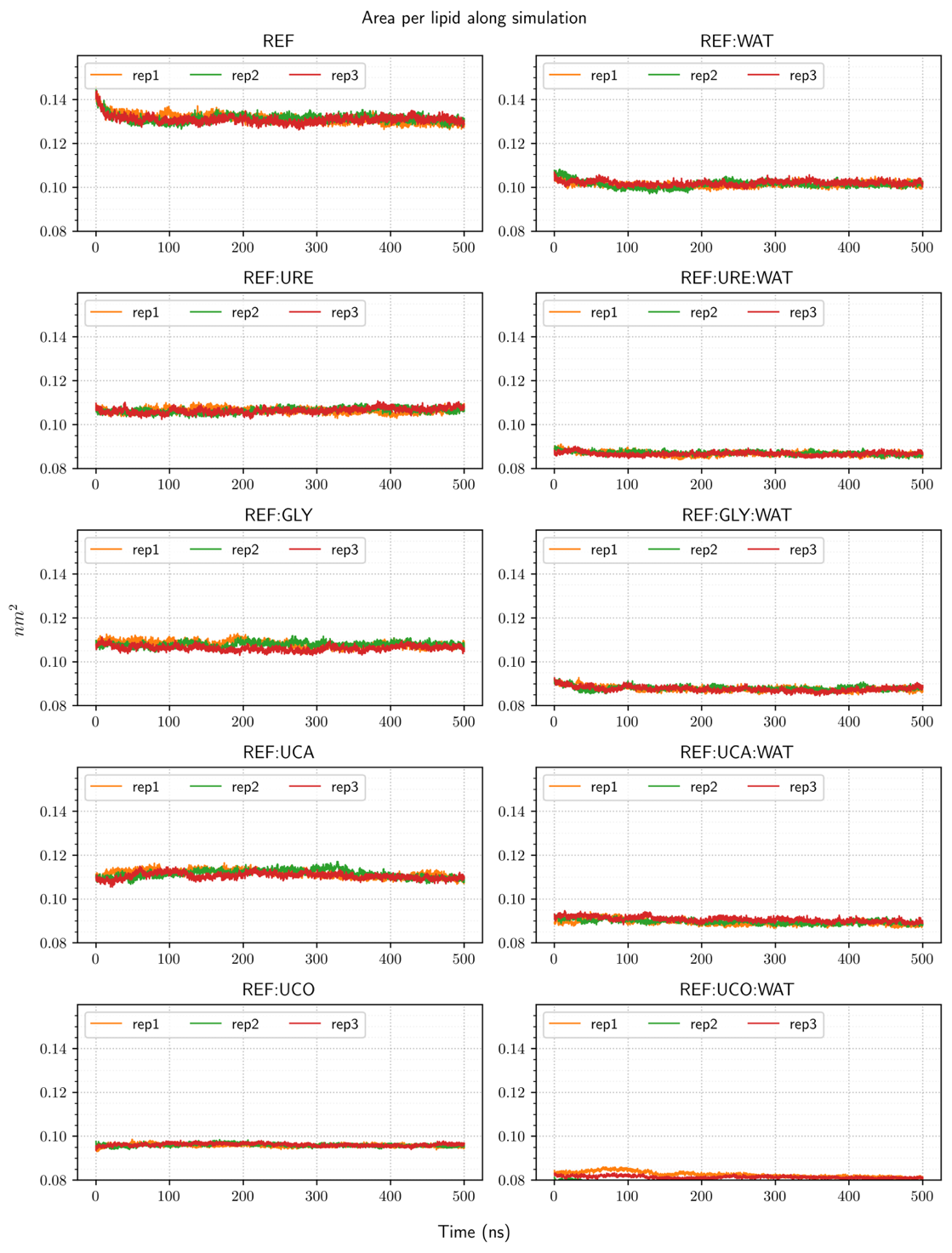

**Figure S1.** Overall area-per-lipid measured along the simulation for each system

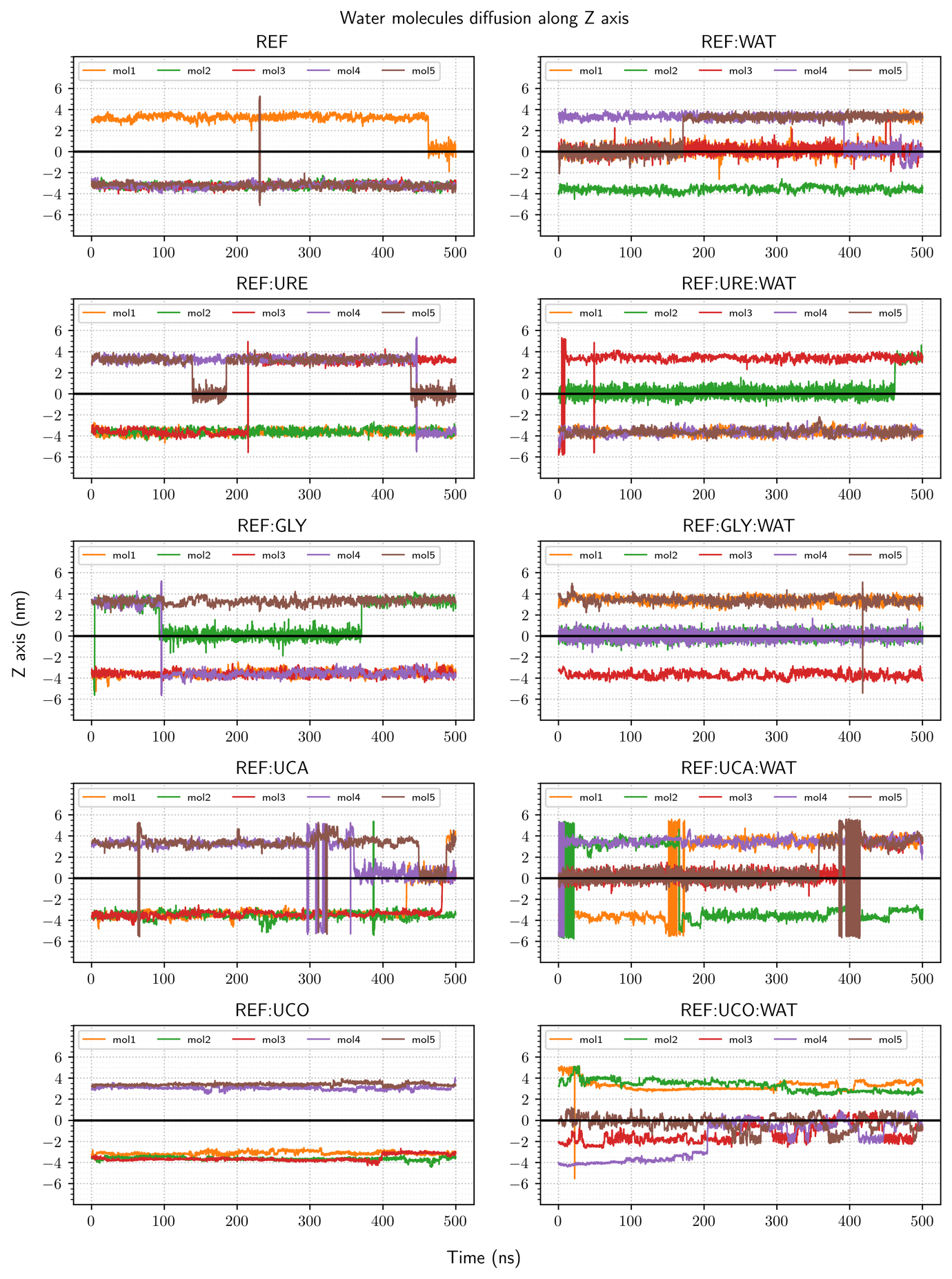

**Figure S2.** Evolution of the position observed along the Z-axis of 5 water molecules randomly chosen and tracked between two hydration levels of each system.

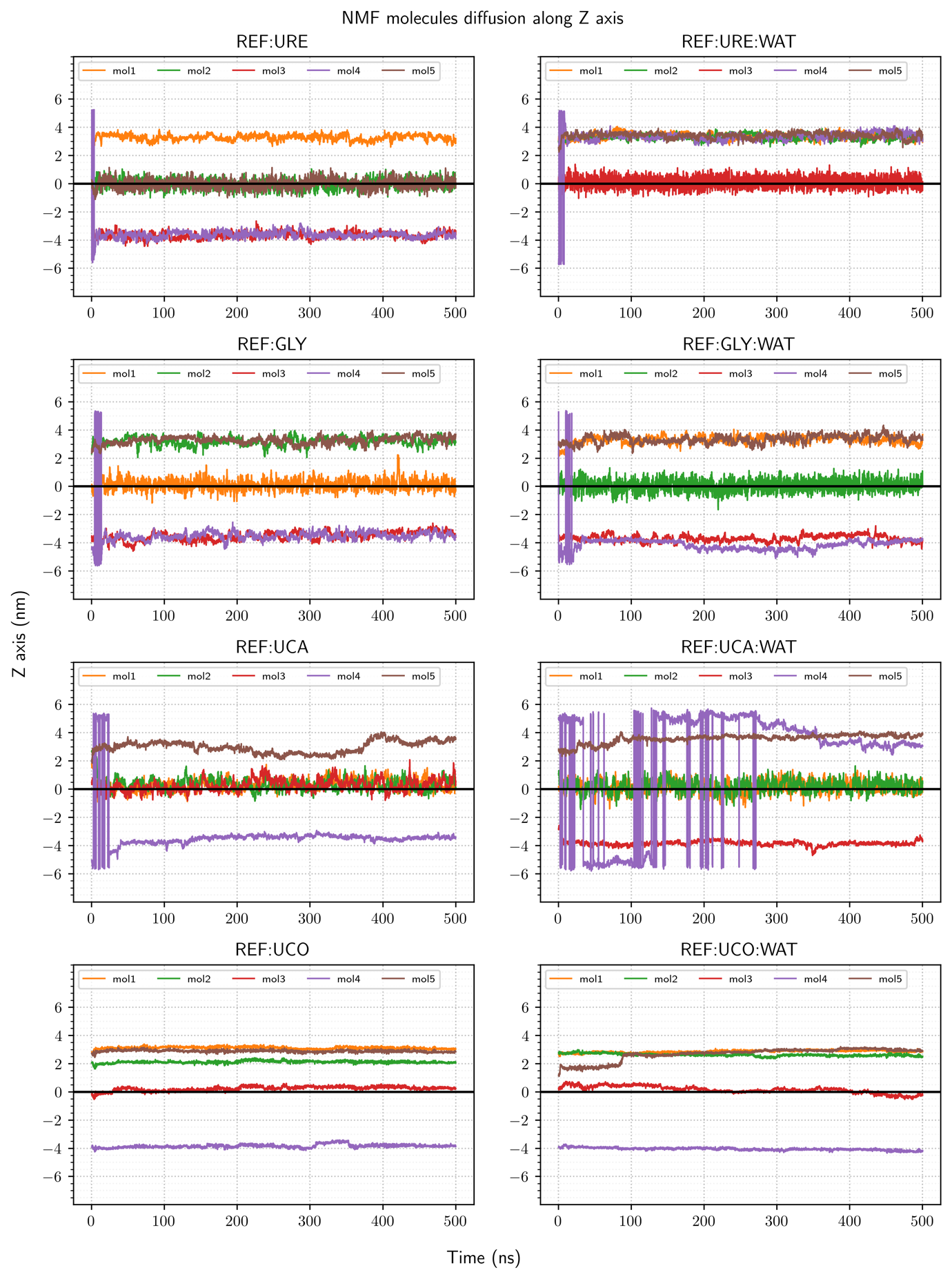

**Figure S3.** Evolution of the position observed along the Z-axis of 5 NMF molecules randomly chosen and tracked between two hydration levels of each system.

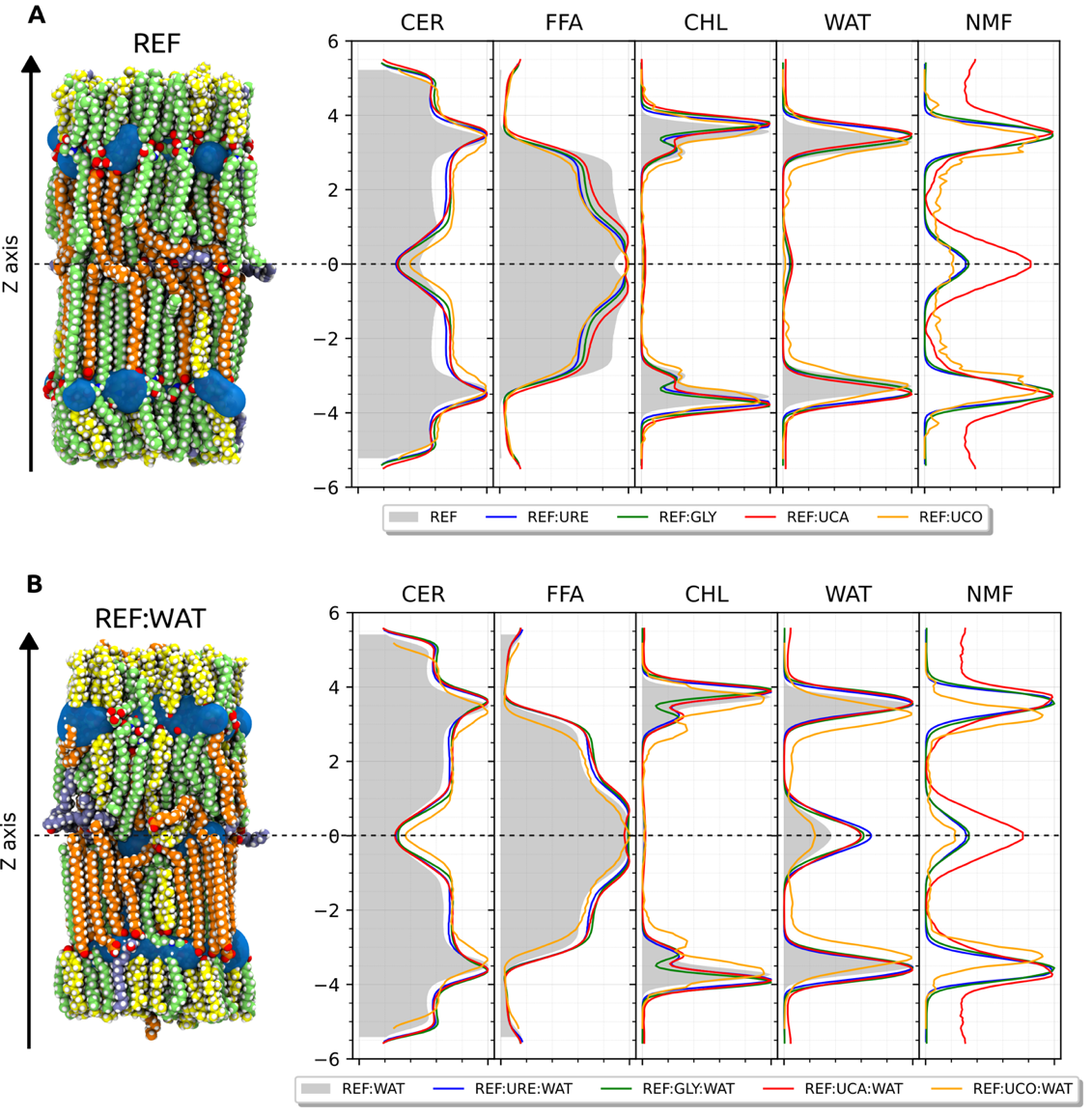

**Figure S4.** Position density occupied by system component along the Z axis. Water (WAT), ceramides (CER), free fatty acid (FFA), cholesterol (CHL), and the corresponding Natural Moisturizing Factor (NMF) of each system.

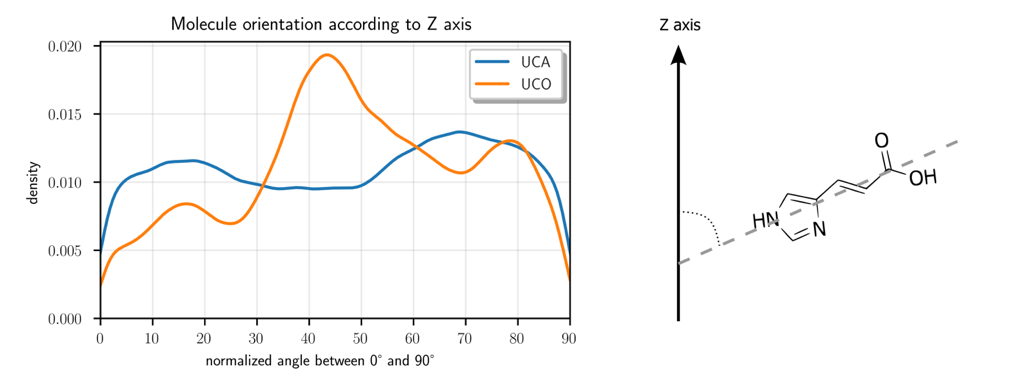

**Figure S5.** UCA and UCO molecules angle orientation according to the z-axis

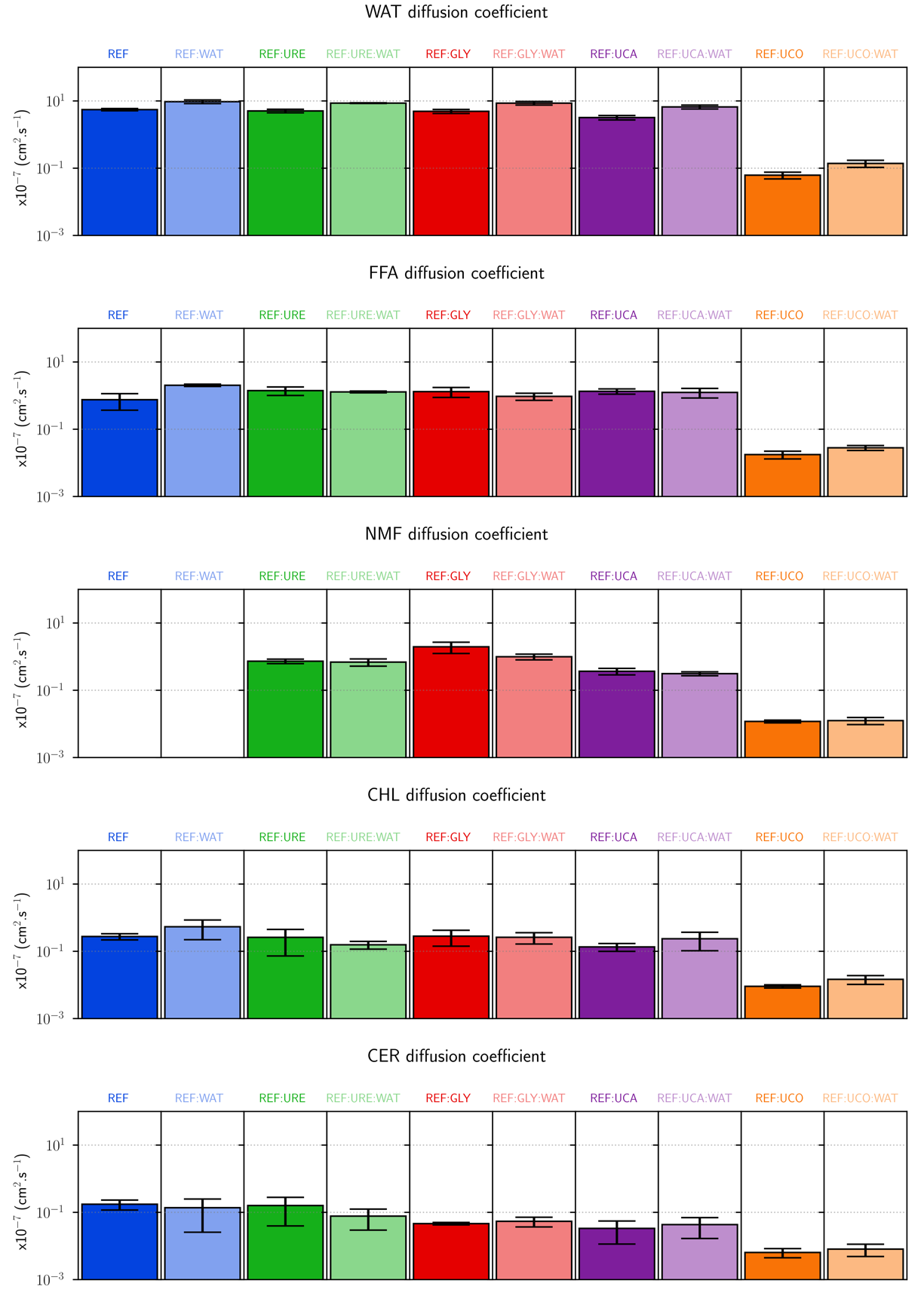

**Figure S6.** Overall diffusion coefficient of each system component

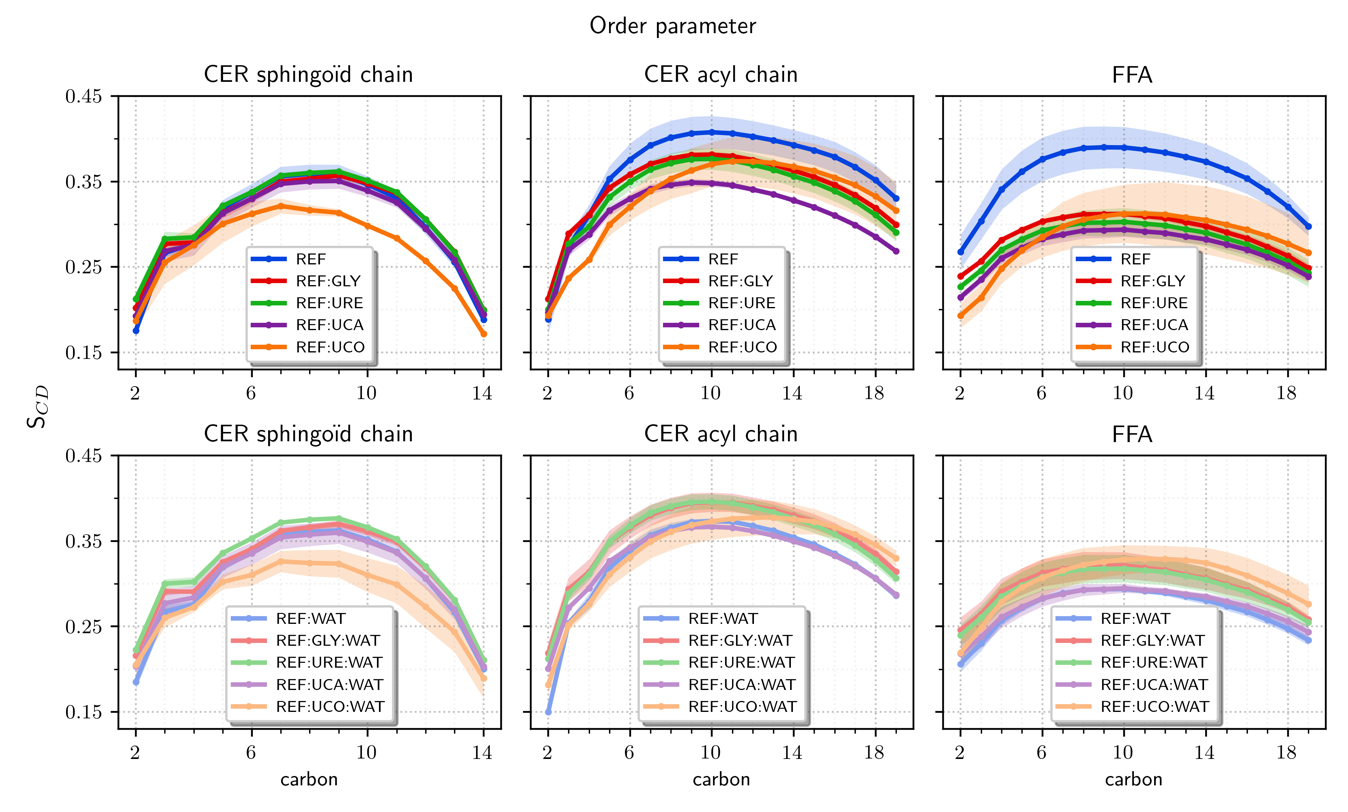

**Figure S7.** FFA and CER chains order parameter. For UCO, consider mainly the first two carbons as this unprotonated form is likely to exist around there, so closer to polar headgroups.

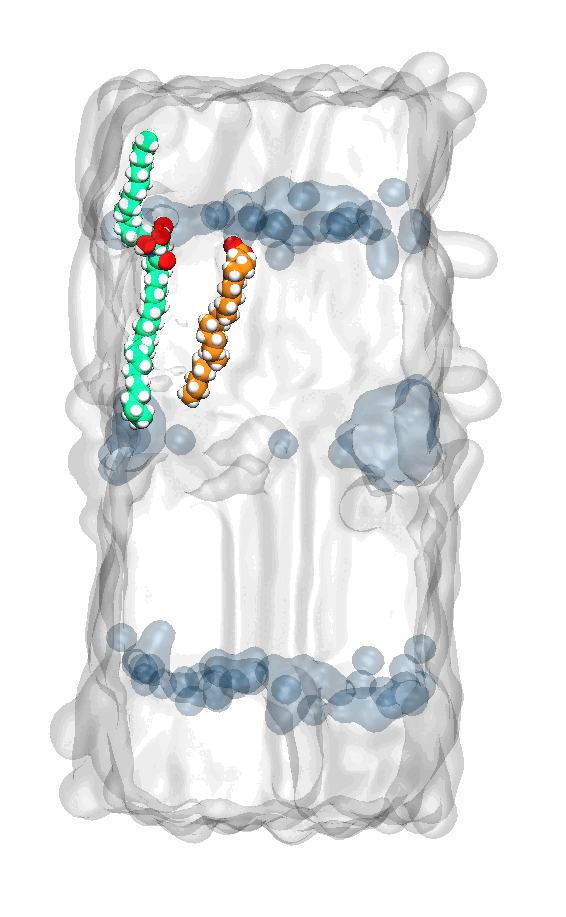

**Figure S8.** Short clip showing the splayed-to-hairpin conformation change of a CER (green) following an FFA (orange) insertion in the short chain layer with water around (blue).
